# Simultaneous targeting of AMPK and mTOR is a novel therapeutic strategy against prostate cancer

**DOI:** 10.1101/2023.08.14.553275

**Authors:** Gangyin Zhao, Gabriel Forn-Cuní, Marvin Scheers, Pier Pieterszoon Lindenbergh, Jie Yin, Quint van Loosen, Leonardo Passarini, Lanpeng Chen, B. Ewa Snaar-Jagalska

## Abstract

Metastatic colonization by circulating cancer cells is a highly inefficient process. To colonize distant organs, disseminating cancer cells must overcome many obstacles in foreign microenvironments, and only a small fraction of them survives this process. How these disseminating cancer cells cope with stress and initiate metastatic process is not fully understood. In this study, we report that the metastatic onset of prostate cancer cells is associated with the dynamic conversion of metabolism signaling pathways governed by the energy sensors AMPK and mTOR. While in circulation in blood flow, the disseminating cancer cells display decreased mTOR and increased AMPK activities that protect them from stress-induced death. However, after metastatic onset, the mTOR-AMPK activities are reversed, enabling mTOR-dependent tumor growth. Suppression of this dynamic conversion by co-targeting of AMPK and mTOR signaling significantly suppresses prostate cancer cell and tumor organoid growth *in vitro* and experimental metastasis *in vivo*, suggesting that this can be a therapeutic approach against metastasizing prostate cancer.

## 1. Introduction

Prostate cancer (PCa) is the second most frequent cancer in males [1]. One of the major causes of death in PCa is incurable bone metastasis, which occurs in 20%-30% of the patients [2–4]. Metastasis is a complex, multiple-step process starting from single cells/clusters that disseminate from the primary site and enter circulation in blood flow until they extravasate and colonize distant organs. While in circulation, the vast majority of the disseminated cancer cells regress due to multiple cell stress factor like anoikis, shear stress, nutrient deficiency, oxidative stress, and immune challenges [5, 6]. Only very few disseminated cancer cells are able to survive the stress and eventually progress into metastases [7–11]. How these cells adapt to the stress and initiate metastatic tumor growth is still elusive due to a lack of tools to trace metastatic seeding of circulating tumor cells *in vivo*.

Chemotherapy targeting specific metabolic pathways in tumor cells has proven to be an effective treatment strategy [12]. One such pathway is the Mammalian Target of Rapamycin (mTOR) signaling cascade. mTOR, a classical protein kinase functions as a central regulator of cell energy sensing, protein synthesis and proliferation. Several inhibitors have been developed to target mTOR, including rapamycin [13], everolimus [14], and temsirolimus [15], among others, all of them receiving FDA approval for the treatment of various diseases. In particular, Everolimus and temsirolimus have been approved for the treatment of several cancers, such as breast cancer, renal cell carcinoma, pancreatic neuroendocrine tumors, and meningioma [16, 17]. However, despite the effectiveness of mTOR inhibition in combating tumor growth, a significant proportion of tumor cells still exhibit resistance to mTOR inhibitors [18–20]. This suggests that relying solely on mTOR inhibitors may not be sufficient to completely eradicate tumors.

Numerous reports have demonstrated that AMP-activated protein kinase (AMPK) can engage in cross-talk with mTOR [21, 22]. AMPK is a heterotrimeric serine/threonine kinase that regulates the levels of ATP, ADP, and AMP in the cell. AMPK is activated by reduced ATP/ADP ratio, for example when the cells are challenged with metabolic and oxidative stress [23]. Once activated, AMPK maintains cellular energy levels by inhibition of ATP-consuming processes such as translation of proteins, synthesis of cholesterol, and fatty acids [24, 25]. Activated AMPK also promotes autophagy, a self-recycling process degrading intracellular components to preserve the levels of ATP during energy deficiency [26, 27]. Additionally, active AMPK inhibits cell proliferation via suppression of mTOR [28–30].

However, despite being first identified as a tumor suppressor due to its inhibitory effect on malignant cell proliferation, which can be related to its mTOR inhibition, a cancer-promoting role of AMPK is emerging. AMPK assumes a crucial role in regulating cellular metabolic balance in situations where mTOR is inhibited [22, 31], and activation of AMPK facilitates the survival of tumor cells during metabolic stress [32]. In breast cancer, for instance, AMPK promotes metastasis by allowing the cancer cells to survive metabolic and oxidative stresses through maintenance of TCA cycle [33]. Similarly, AMPK protects leukemic stem cells from metabolic stress in bone marrow [34]. In prostate cancer, the inhibitory effect of AMPK on cancer development has been well documented [35], but it is still unclear if AMPK can protect prostate cancer cells in certain stressful conditions, like in circulation and seeding at foreign microenvironment, or during mTOR chemical inhibition.

In summary, while mTOR inhibitors have shown promise as a targeted chemotherapy approach, resistance to these inhibitors remains a challenge. The intricate relationship between mTOR and AMPK highlights the complexity of tumor metabolism and the need for comprehensive treatment strategies to effectively combat tumors. In this study, we used FRET-based AMPKα kinase biosensor, zebrafish xenografts, and PDX-derived organoids and found a dynamic reversal of mTOR and AMPK signaling in prostate cancer cells during metastasis. This conversion enables cancer cells to adapt to metabolic stress when circulating in blood flow, and initiate tumor growth after metastatic seeding. Suppression of this conversion by co-targeting of AMPK and mTOR signaling significantly suppresses experimental metastasis onset and growth of PDX-derived organoids.

## 2. Materials and Methods

### 2.1. Cell-line maintenance and cell line construction

Prostate cancer cell lines PC3 and PC-3M-Pro4 were cultured in DMEM with 10% FCS (Fetal Bovine Serum) and were split 1:5 every 3-4 days. For 3D culturing, cells were seeded in Corning 96-well CellBIND microplates (Sigma Aldrich, cat#: CLS3340) and media and treatments were refreshed at day 1, 4 and 8.

Short hairpin RNA (shRNA) constructs were obtained from Sigma’s MISSION library (kindly provided by Department of Molecular Cell Biology, LUMC). Other plasmids were purchased from the Addgene library: pLenti-Utra 6.3-dtomato (#106173), pLenti-GFP-puro (#17481), pLenti: Fli:GFP-T2A-mCherry [36] (#124428), pLenti-constitutively active AMPK (#139843), and pLenti-H2B-GFP (#51004). The pLenti-AMPK Biosensor was a kind gift from Sylvia Le Dévédec (Addgene plasmid #50526). Lentiviral particles were produced by transforming pLenti constructs, packaging plasmids psPAX2, and envelop plasmids pMD2.G (a gift from Dr. Maciej Olszewski) into HEK-293T cells using lipoD293 (SignaGen Laboratories) as transforming reagent. Lentivirus present on the supernatant were collected 72 hours after transformation. Cells were transduced with the produced lentivirus using 6 μg/ml Polybrene (Sigma-Aldrich) and selected using media containing 10 μg/mL blasticidin or 2 μg/mL puromycin according to the resistance sequence carried by the lentivirus.

### 2.2. Zebrafish tumor xenograft model

Zebrafish were handled in compliance with the local animal welfare directives of Leiden University (License number AVD1060020172410 and AVD10600202216495) and following standard zebrafish rearing protocols (https://zfin.org), which adhere to the international guidelines from the EU Animal Protection Direction 2010/63/EU. Lines used in this study were the transgene fish lines Tg(mitfa^w2/w2^; mpv17^a9/a9^), also known as Casper, and Tg(mitfa^w2/w2^; mpv17^a9/a9^; y1Tg) (ZfinID ZDB-FISH-180529-3), from here on mentioned as Tg(fli:EGFP)/Casper. Embryos were collected and maintained in egg water (Sera Marin salt, 05420; 60 µg/ml in distilled deionized water) at 28°C for the first 2 days post fertilization (dpf).

At 2 dpf, hatched zebrafish embryos were engrafted via the Duct of Cuvier with 1×10^5^ cells/µL tumor cells resuspended in 2% polyvinypyrrolidone 40 (PVP-40, Calbiochem, San Diego, USA) using borosilicate glass capillary needles (1mm O.D. x 0,78mm I.D., Harvard Apparatus) pulled with Model P-97 needle puller (Sutter Instrument Company). After engraftment, larvae were kept at 33°C with daily egg water refresh for a maximum of 6 days post-injection (dpi), when fish were euthanized and all experiments ended. Zebrafish larvae were screened at 1 dpi for successful grafting and no edema. Prior to injections and imaging, embryos were anaesthetized with 0.02% buffered 3-aminobenzoic acid ethyl ester (tricaine; Sigma-Aldrich, A-5040) in egg water.

Treatment with SBI-0206965 (ApexBio: #A8715) and/or Rapalink-1 (ApexBio: #A8764) was administrated by IV injection 4 hours after xenografting, followed by a second administration 18 hours later. Unless otherwise specified, 40-50 fish were used per experimental group.

### 2.3. Western blot

At 1, 2, 3, 4, 5, and 6dpi, 150 tails of zebrafish larvae xenografted with PC-3M-Pro4 were collected and digested with 1 μg/ml Liberase TL at 34°C for 20 min, mixed every 2 min. The reaction was stopped with 10% FCS and centrifuged at 1000rpm at 4°C for 5 min. After removal of supernatant, cells were resuspended with 5 ml PBS and filtered with a 70 μm filter membrane. Total protein was purified by the AllPrep DNA/RNA/Protein Mini Kit (Qiagen: # 80004). Corresponding protein expression levels in the cells of different groups were detected by western blot using the following antibodies: p-AMPK (Cell signalling: #2535S and #2532S), AMPK (Santa Cruz Bio: sc-74461), mTOR (Santa Cruz Bio: SC-517464), AKT (Cell signaling: #9272), p-AKT (Cell Signaling: #4060), p70S6k (Cell signaling: #9202), p-p70S6k (Cell Signaling: #9204), p-ACC (Santa Cruz Bio: sc-271965), and ACC (Cell Signaling: #3676). Bands were visualized by the max clarity Western ECL substrate (Bio-Rad) on the Chemidoc Universal Hood III (Bio-Rad).

### 2.4. Cell viability and death detection by WST-1 and LDH

Cell proliferation and cell death were assessed using the WST-1 and cytotoxicity Detection KitPLUS (LDH) (Roche) following the manufacturer’s protocol. Briefly, 5000 cells in 100 µL of DMEM with 10% or 2.5% FCS were seeded in each well of a 96-well plate. After 24 h, the medium was renewed with medium containing either Vehicle (Veh) control or the specified inhibitors. Cells were allowed to grow for 72 h. For cell viability measurement, 10 µL/well WST-1 reagent was added followed by 2 h incubation at 37 °C before absorbance reading using a microplate reader M1000 PRO (TECAN). Each condition was repeated 6 times in a minimum of two independent experiments.

### 2.5. Colony assay, invasiveness assay and cell proliferation detection

Colony assay was performed using crystal violet staining [37]. For this, 2000 cells were seeded in a 6-well plate and grown for a maximum of 2 weeks. Once wells were confluent, cells were washed twice with ice-cold PBS. Next, the cells were fixed by 500 µL ice-cold 100% methanol and incubated on ice for 10 minutes, followed by removal of the methanol and addition of 500 µL of a 50% mix of glycerol and PBS into each well. Staining was performed by removing the glycerol/PBS mix and covering the cells with crystal violet at room temperature for 10 minutes. Images for quantification were taken after three washing steps.

3D collagen invasion assays were performed using 1 mg/ml Collagen type I stock (BD Biosciences: #354265) in 44 mM NaHCO3, 0,1M HEPES, and DMEM. 1 mL collagen was added per well of a 6-well plate and solidified in at 37°C. Cancer cells were loaded into a Borosilicate glass capillary needle (1mm O.D. x 0,78mm I.D., Harvard Apparatus) and injected into the collagen using a microinjector with the PV820 Pneumatic PicoPump (WPI). Approximately 500 cells were injected and imaged at days 0, 1 and 2 after injection. Changes in cell proliferation were determined using the WST-1 assay kit (Abcam).

### 2.6. Immunohistochemistry

2D cultured cells in 8 wells chamber were fixed with 4% PFA (Sigma-Aldrich) in PBS and permeabilized with 0.5% triton-X 100 in PBS. After incubation with blocking buffer containing 5% BSA in PBS, cells were incubated with primary antibody (rabbit Ki67; Abcam: ab16667) overnight at 4°C and with fluorescent conjugated secondary antibody, and DAPI (Thermo Fisher) for 1 hour at room temperature before imaging. Whole-mount immunostaining of zebrafish and LAPC9 organoids was performed as previously published [38–40].

### 2.7. Organoid culture and testing

LAPC9 organoids were a kind gift from Marianna Kruithof-de Julio and cultured using published methods [40, 41]. For drug pre-treatment, LAPC9 organoids were cultured in 96-well low attachment plates in complete PCa medium for 72 h, after which the medium was replaced with fresh medium containing drugs. Organoids were cultured for another 5 days before proceeding with microscopy image and viability test by CellTiter-Glo 3D kit (Promega: G9681) following manufacturer’s instructions.

### 2.8. Imaging, quantification, data analysis and statistics

Images were taken with a M205 FA stereo microscope (Leica) or an SP8 confocal microscope (Leica) using a 40X immersion objective (NA = 1.4) and equipped with 488-nm, 532-nm, and 638-nm laser lines. Dose-response curves, matrixes, and Bliss Synergy scores were calculated and plotted using SynergyFinder 3.0 [42]. Image-based quantifications and representative images were created using Fiji [43]. Data analysis and statistics were performed using either ImageJ (Fiji) or Graphpad Prism version 9. Unless otherwise stated, all experiments were repeated up to a total of three independent replicates. Significant results were noted as follows: *P-value = <0,5, **P-value = 0,01, ***P-value = 0,001 & ****P-value = 0,0001.

## 3. Results

### 3.1. Metastatic onset is associated with conversion of metabolism signaling pathways governed by AMPK and mTOR

In order to study metastatic onset of prostate circulating tumor cells at a single cell resolution, we used a previously reported xenograft zebrafish model [44] and intravenously (IV) injected mCherry-labeled PCa cells (PC-3 and PC-3M-Pro4). After injection, cancer cells circulated in blood flow until 1dpi, adhered to endothelial cells (1dpi – 2 dpi), extravasated (2dpi – 3dpi), and grew into perivascular micro-metastases at the caudal hematopoietic tissue (CHT at 4dpi – 6dpi), the major hematopoietic organ in zebrafish larvae at this stage (**Fig. 1A, S1A, S1B, Supplementary Video 1**).

**Fig. 1.**
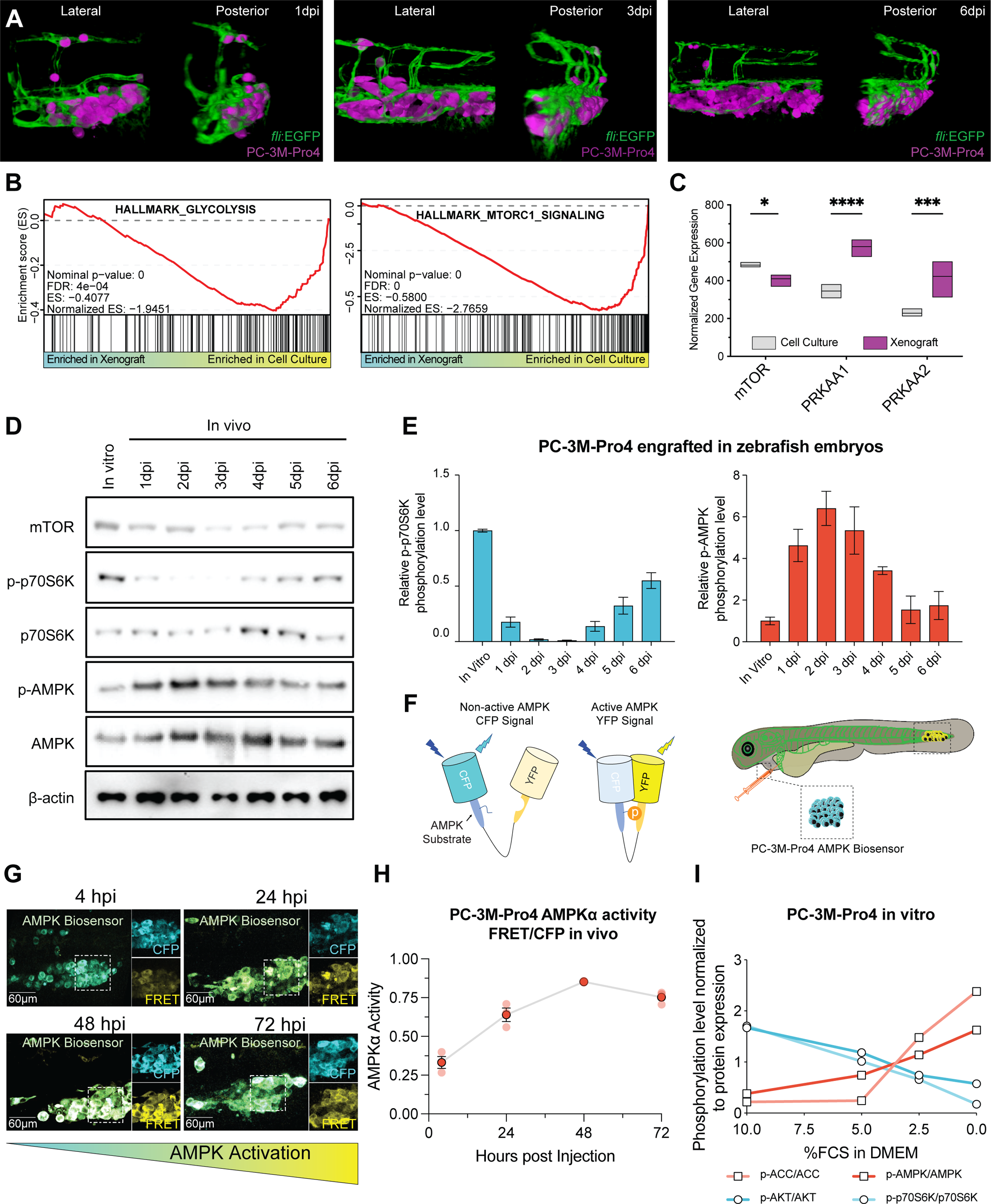
The disseminating prostate cancer cells display decreased mTOR and increased AMPK activities. **A.** High resolution 3D image at 40X magnification of the metastatic foci in zebrafish embryos at different days after engraftment. PC-3M-Pro4 cells expressing tdTomato are labelled in magenta, while blood vessels expressing GFP are shown in green. **B.** Gene Set Enrichment Analysis results indicate that genes related to glycolysis (left) and mTOR pathway (right) were enriched in the cell culture compared to engrafted cells. **C.** Normalized transcriptomic gene counts show decreased expression for mTOR, and increased expression AMPKα1 (PRKAA1) and AMPKα2 (PRKAA2) in prostate cancer cells at 6 dpi in zebrafish xenograft vs cell culture. **D, E**. Western blot of phosphorylated proteins from the AMPK and mTOR pathways indicates higher AMPK activity and lower mTOR activity during the experimental timeline. **F**. Schematic representation of the PC-3M-Pro4 AMPK biosensor cell line injected in zebrafish in this study. **G.** Representative confocal image of PC-3M-Pro4 AMPK-biosensor after engraftment. Blue represents CFP signal, yellow represents FRET signal and activation of AMPK. **H**. Quantification of the PC-3M-Pro4 AMPK activity during the first 72 hours post engraftment (n = 5) shows activation of AMPK during the first hours post engraftment. **I**. Phosphorylation levels of proteins downstream of activation of AMPK and mTOR pathways demonstrate high AMPK activity and low mTOR activity during metabolic stress in prostate cancer cells.

We first reanalyzed a previous transcriptomic study to examine how the injected cancer cells responded to the microenvironment at the metastatic onset by comparing the transcriptome of PC-3M-Pro4 at the metastatic foci in xenografts (6 dpi) with those before engraftment (**Fig. S1C**) [45]. Transcriptional programs indicative of metabolic stress such as ATP metabolic process, positive regulation of autophagy, cellular response to glucose starvation, and cell cycle arrest were among the most positively enriched gene signatures at the metastatic onset (**Fig. S1D)**. In contrast, gene signatures for carbon metabolism like glycolysis, TCA cycle, and mTOR signaling pathway were negatively enriched (**Fig. 1B, S1D**). This transcriptomic analysis strongly suggested that engrafted cancer cells experienced metabolic stress during metastatic initiation. Congruent with this observation, genes encoding for the energy sensor AMPK subunits AMPKα1 (PRKAA1) and AMPKα2 (PRKAA2) were significantly upregulated at metastatic onset while transcription of mTOR was downregulated (**Fig. 1C**). We then confirmed this AMPK-mTOR signaling conversion over time after engraftment via western blot (Supplementary **Fig. S1E**). Phosphorylation of the mTOR downstream target p70S6K was significantly reduced in the first 3dpi and then gradually recovered at 6dpi. In contrast, phosphorylation of AMPK (correlated with its activation) was elevated compared to their *in vitro* controls, with a peak at 2dpi followed by a gradual decrease until 6dpi (**Fig. 1D, 1E)**. We confirmed the kinetic activity of AMPK in the engrafted cancer cells using a FRET-based AMPKα kinase biosensor, a fluorescent indicator for AMPK activity in living cells (**Fig. 1F**) [46]. Consistent with the protein analysis, FRET signal resembling AMPK activation was continuously increased in the cancer cells at first 3 days after transplantation (**Fig. 1G, 1H)**. Overall, these data strongly demonstrate that the metastatic onset of prostate cancer cells is associated with a transient activation of AMPK and deactivation of mTOR signaling pathway.

We hypothesized that the mTOR-AMPK signaling conversion could be mediated by metabolic stress. To mimic the stress conditions that cancer cells endure during circulation *in vivo*, we cultured them in medium supplemented with a reduced concentration of fetal calf serum (FCS). Consistently, we found that AMPK and its downstream target ACC were increasingly phosphorylated, while the mTOR signaling components AKT and p70S6K were dephosphorylated while lowering FCS concentrations (**Fig. 1I, S2A)**. Moreover, this signaling conversion led by serum starvation was accompanied by cell cycle arrest as defined by reduced Ki67 expression (**Fig. S2B, S2C**). These data confirm that the mTOR-AMPK conversion at the metastatic onset could indeed be caused by metabolic stress.

### 3.2. AMPK protects prostate cancer cells from cell death in vitro and in vivo

Having shown the mTOR-AMPK conversion in the cancer cells at early stage of metastasis, we hypothesized that this signaling conversion is required to protect the cells from metabolic stress-induced death. To investigate this, we firstly used short hairpin RNA (shRNA) to knock down both AMPK subunits AMPKα1 and AMPKα2 (Dkd) in PC-3M-Pro4 and PC-3 cell lines (**Fig. S3A**). We then determined the effects of the AMPKα Dkd on cell survival and growth compared with control shRNA (SCR) by a clonogenicity assay. Double knockdown of AMPK slightly impaired long-term culture growth in a rich nutrient environment (10% FCS), but the inhibition was significantly stronger for the cells in the nutrient depleted medium with 2.5% FCS (**Fig. S3B, S3C)**. Similarly, viability of the AMPKα Dkd cells in nutrient deficiency was dramatically decreased compared to the SCR control cells (**Fig. S3-I**), confirming that AMPK plays an important protective role in prostate cancer cell survival under metabolic stress.

Next, we interrogated the importance of AMPK during metastatic onset by engrafting SCR or AMPKα Dkd PC-3M-Pro4 and PC-3 cells and measuring tumor growth at the CHT of zebrafish embryo using fluorescent microscopy. Cancer cells with AMPKα Dkd were significantly impaired in their ability to grow a metastatic tumor at 3dpi and 6dpi (**Fig. 2A-D**). We hypothesized that the AMPK signaling is required to protect the cancer cells from stress-induced death in the microenvironment. We therefore used immunofluorescence to measure cleaved Caspase 3, a marker of apoptosis, in the cancer cells at an early time point post engraftment (2dpi) (**Fig. S3J**). The signal for cleaved Caspase 3 was significantly increased in the AMPKα Dkd cells compared to the control at the metastatic site, demonstrating AMPK plays a protective role in the survival of disseminating cancer cells (**Fig. 2E-H, S3K, S3L**).

**Figure 2.**
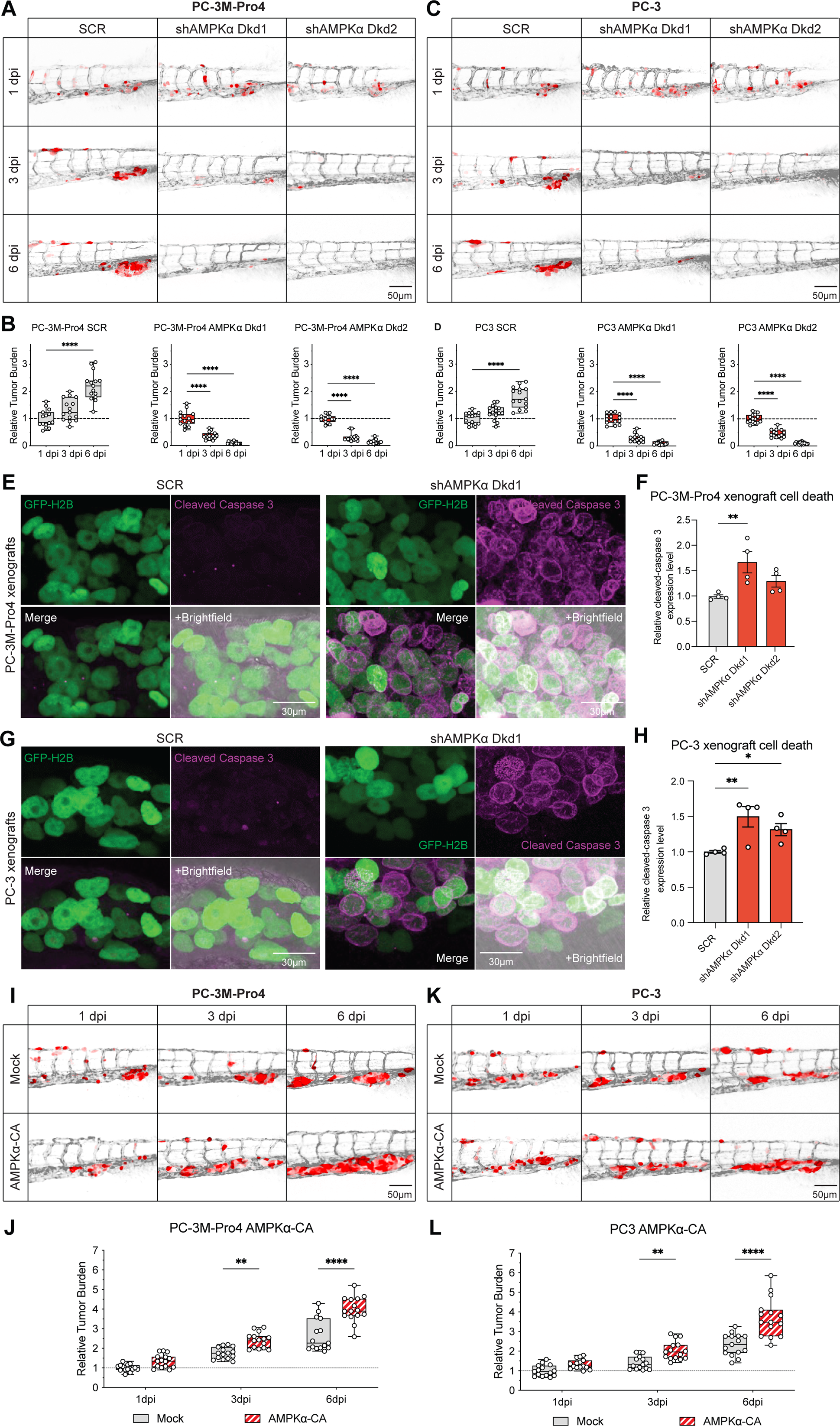
AMPK knockdown induces tumor cell death while its activation induces cell survival in circulation. **A.** Representative images of the metastatic foci at 1, 3, and 6 dpi in zebrafish embryos xenografted with PC-3M-Pro4 expressing tdTomato and transduced with control (SCR) or two replicates of a double AMPKα1 and AMPKα2 knockdown (shAMPKα Dkd). **B.** The shAMPKα Dkd PC-3M-Pro4 cells show reduced ability to form metastasis in zebrafish embryos (n = 15). For relative tumor burden quantification, integrated fluorescence intensity was measured using Fiji. **C, D.** The same results were obtained using the PC-3 cell line background. **E.** Representative images of immunostaining of the cell nucleus (GFP-H2B, green) and apoptosis (cleaved-caspase 3, magenta) in PC-3M-Pro4 SCR and shAMPKα Dkd xenografted in zebrafish embryos. **F.** Quantification of the relative cleaved-caspase 3 signal showes that shAMPKα Dkd cells undergo higher cell death rate, n = 3. **G, H.** The same results were obtained using the PC-3 cell line background. **I.** Representative images of the metastatic foci of PC-3M-Pro4 expressing tdTomato with or without constitutively activated AMPK (AMPKa-CA) at 1, 3, and 6 dpi. **J.** The PC-3M-Pro4 cells expressing AMPKa-CA show increased ability to form metastasis in zebrafish embryos. **K, L.** The same results were obtained using the PC-3 cell line background.

To further confirm the role of the active AMPK in cancer cell survival and growth at early stage of metastasis, we overexpressed constitutively activated AMPK (AMPKa-CA) [47] in PC-3M-Pro4 and PC3, and engrafted those cells into zebrafish. AMPK overexpression significantly increased survival and growth of the injected cancer cells at the metastatic site (**Fig. 2I-L**). Taken together, these data strongly support that active AMPK is required to protect the disseminated cancer cells from stress-induced cell death during metastatic onset.

### 3.3. AMPK is activated and promotes cell survival during mTOR inactivation

The indispensable role of the active AMPK in prostate cancer metastasis prompted us to further interrogate mechanisms of AMPK regulation in prostate cells during metastatic onset. Given that we had previously seen that AMPK activation and mTOR deactivation were closely associated during that time, we questioned if AMPK activity could be directly triggered by the stress-induced deactivation of mTOR signaling. We therefore treated the cancer cells with the third-generation dual mTORC1/2 inhibitor Rapalink-1 [48, 49], which significantly suppressed phosphorylation of the mTOR signaling components AKT and p70S6K. The phosphorylation of AMPK and ACC, on the other hand, was increased, indicating AMPK signaling was activated by this treatment (**Fig. 3A, 3B**). We further demonstrated AMPK activation following mTOR inhibition using a FRET-based AMPK assay: the AMPK FRET signal was significantly increased in the cells treated with Rapalink-1 (**Fig. 3C, 3D** and **Supplementary video 2**).

**Figure 3.**
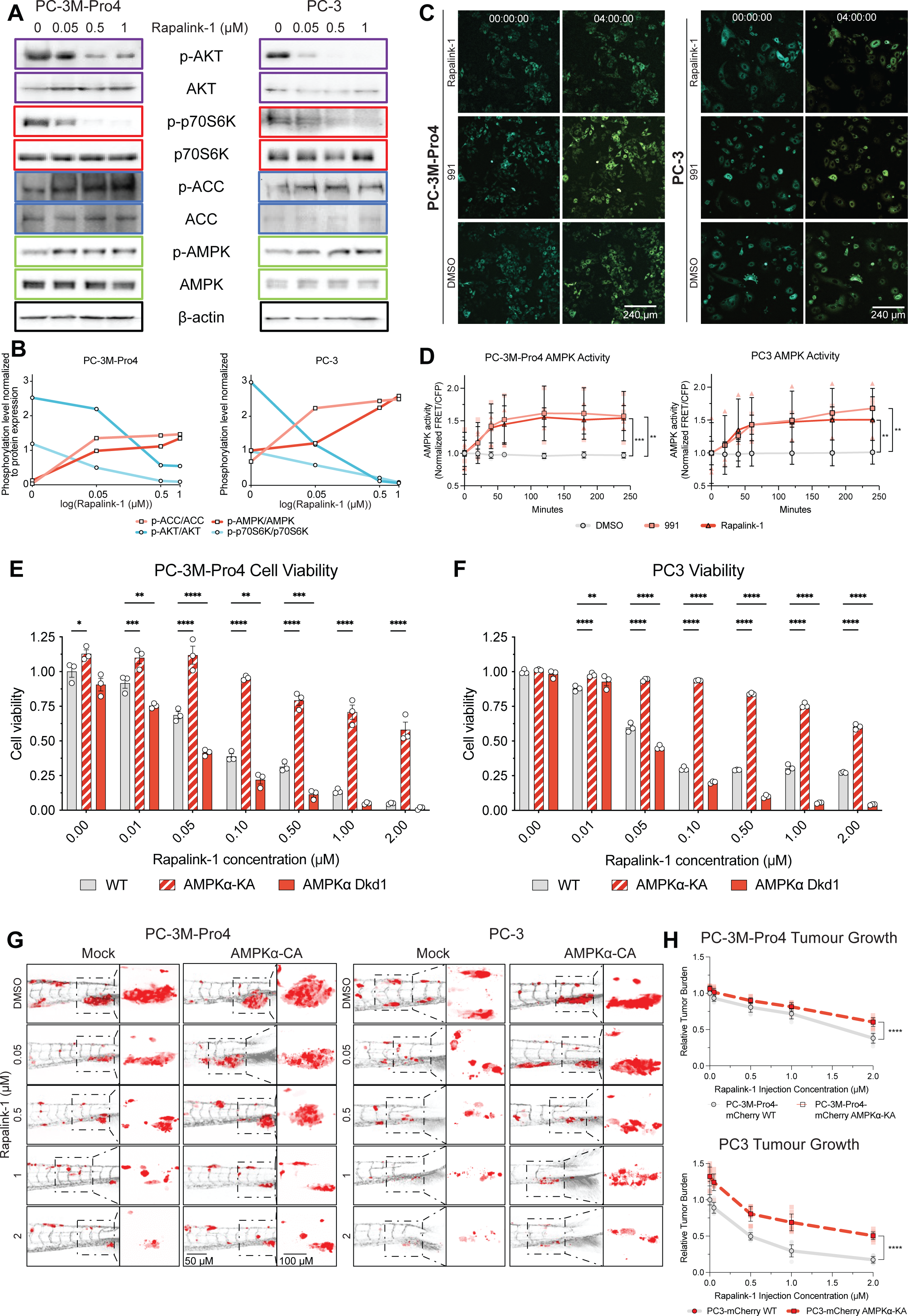
Inhibition of mTOR activates AMPK, making cells partially resistant to Rapalink-1. **A, B.** Western blot of phosphorylation levels of proteins downstream of mTOR and AMPK activation demonstrated that Rapalink-1 inhibites mTOR and activates AMPK in PC-3M-Pro4 and PC-3 cells in a concentration-dependent manner. **C.** Representative images of PC-3M-Pro4 and PC-3 cells expressing an AMPK biosensor treated with DMSO (negative control), Rapalink-1 (0.5μM), or the small molecule compound 991 (10 μM) (AMPK activator, positive control) at 1 and 3 hours post treatment. **D**. Rapalink-1 activates AMPK in cells as calculated as a ratio of FRET/CFP. **E, F**. PC-3M-Pro4 and PC-3 cells overexpressing constitutively active AMPK (AMPKa-CA) showed stronger resistance to Rapalink-1, while the shAMPKα Dkd cell lines were more sensitive to the drug. **G.** Representative images of the metastatic foci of PC-3M-Pro4 and PC3 expressing tdTomato and overexpressing AMPKa-CA xenografted in zebrafish and treated with different concentrations of Rapalink-1. **H**. The AMPKa-CA overexpressing cells were more resistant to Rapalink-1 induced cell death in vivo (15 fishes/group).

Next, we examined the biological relevance of AMPK activation in the context of mTOR deficiency. Prostate cancer cells bearing either SCR control, AMPKa-CA overexpression or AMPKα Dkd were treated with Rapalink-1 *in vitro*. Compared to the SCR cells, mTOR inhibition had a strong reduction on AMPKα Dkd cell viability, while AMPKa-CA cells were resistant to this treatment (**Fig. 3E, 3F)**. We also tested if AMPK could protect the cancer cells from mTOR deficiency *in vivo*. To do so, cancer cells bearing either mock control or AMPKa-CA were transplanted into zebrafish followed by treatment with Rapalink-1. At nontoxic doses (**Fig. S4A**), the metastatic growth of cancer cells was significantly inhibited in a dose-dependent manner (**Fig. 3G, 3H**). Importantly, consistent with *in vitro* observations, AMPKa-CA cells in zebrafish were significantly more resistant to the treatment. Collectively, these data demonstrate that AMPK is directly activated in prostate cancer cells following inhibition of mTOR, and that this activation can promote prostate cancer cell survival during mTOR inhibition.

### 3.4. Dual inhibition of mTOR and AMPK show synergetic effects against prostate cancer cell viability in vitro

Hyper-activation of mTOR signaling has been identified as one of the major drivers of prostate cancer tumorigenesis [50, 51]. Therefore, targeting of this signaling pathway using mTOR inhibitors has long been considered as a potential approach to treat this disease. Nevertheless, patient responses to the treatment remain poor, mainly due to acquired resistance to the treatment [51, 52]. Following our findings that active AMPK can promote survival of prostate cancer cells during mTOR inhibition, we intended to decipher whether the inactivation of AMPK can pose a vulnerability to prostate cancer cells in the context of mTOR inhibition. Thus, we tested anti-cancer potential of the AMPK inhibitor SBI-0206965 (SBI) [53, 54], in combination with the mTOR inhibitor Rapalink-1.

When cultured cancer cells in monolayer were co-treated with both compounds, viability of the cells was significantly reduced compared to the PC-3M-Pro4 (**Fig. 4A, S5A, S5B**) and PC-3 (**Fig. S5C-5E**) treated with either SBI or Rapalink-1 alone. For PC-3M-Pro4, the highest percentages of inhibition for SBI (10 μM) and Rapalink-1 (0.5 μM) were 28.18% ± 3.34 and 64.48% ± 0.94, respectively, while its combination achieved a 94.42% ± 0.69 of growth inhibition. As shown via the Bliss synergy score analysis, the combination of SBI and Rapalink-1 had strong synergistic effects at middle concentrations for PC-3M-Pro4 (SBI+Rapalink-1 range: 1-10 μM + 0.01-0.5 μM) and relative high concentration for PC-3 (SBI+Rapalink-1 range: 5-10 μM + 0.5-1 μM) (**Fig. 4B** and **Fig. S5F**). Additionally, we evaluated the efficacy of the treatments on 3D tumor spheroids. In order to precisely measure effects of the treatments on cell apoptosis in spheroids, we transduced the cells with the FlipGFP-based caspase reporter for imaging apoptosis [36]. After transduction of the reporter, cancer cells were cultured in ultra-low attachment surface to form tumor spheroids, and then treated with SBI, Rapalink-1 or its combination at serial concentrations for 12 days (**Fig. S6A**). Compared to the SBI or Rapalink-1 single treatments, which only induced mild apoptosis at high doses, the combined treatment significantly increased cell death in the spheroids at lower concentrations (SBI+Rapalink-1 range: 5-10 μM + 0.5-2 μM) (**Fig. 4C, 4D, S6B, S6C**). By calculating the dynamics of apoptosis rate of 3D cells for 12 consecutive days, we demonstrated that the combination of two drugs had a stronger effect on tumor inhibition than mono treatments (**Fig. 4E**). In addition to the induction of apoptosis, the size of the spheroids was also significantly reduced by the co-treatments (**Fig. 4F)**. Altogether, these data indicate that the combination of mTOR and AMPK inhibitors can exert a strong inhibitory effect on prostate cancer cells *in vitro*.

**Figure 4.**
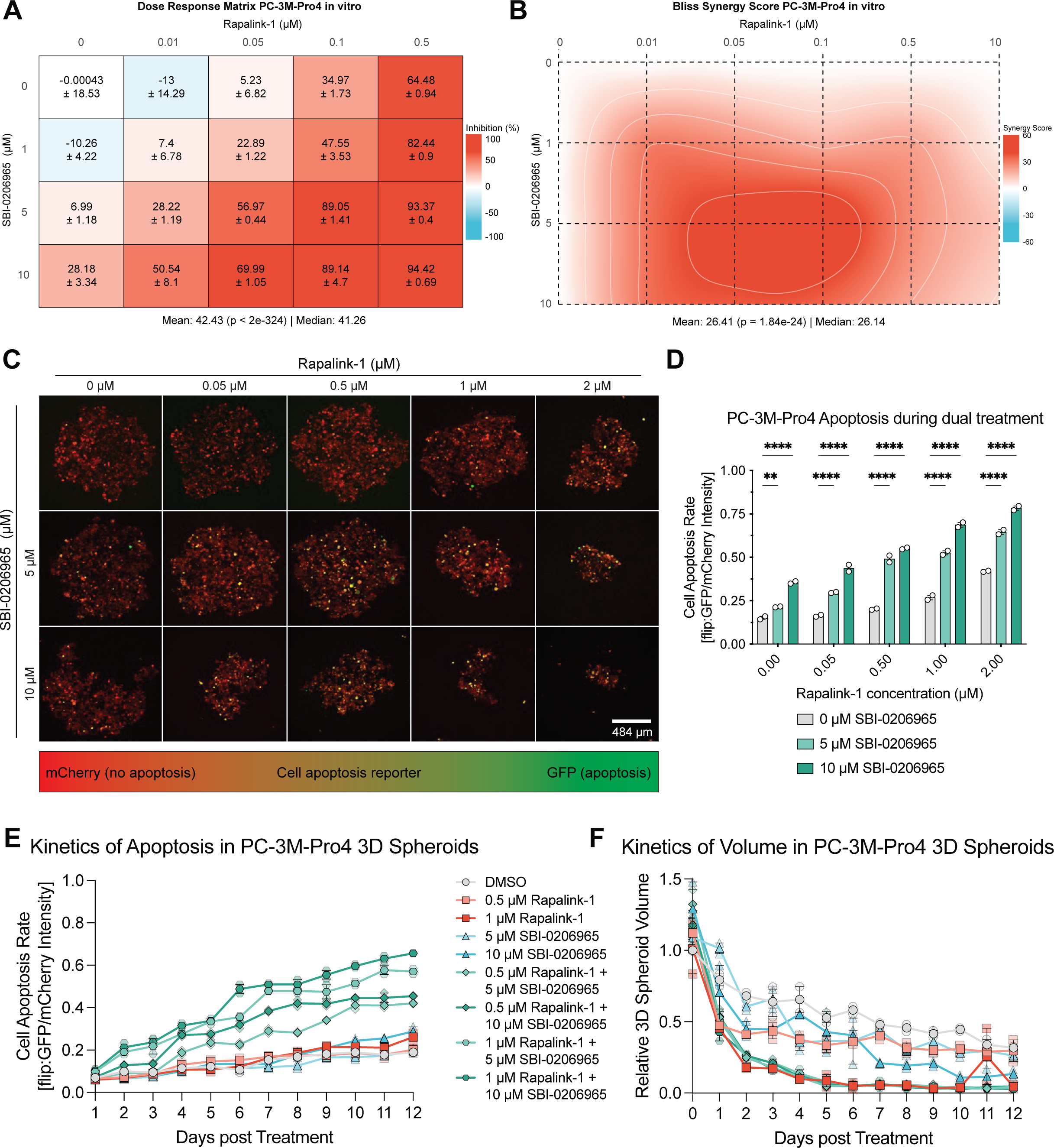
Synergistic use of mTOR and AMPK inhibitors is a highly effective tumor suppressor in 2D and 3D cultures. **A.** Dose-Response matrix of cell growth inhibition of different Rapalink-1 and SBI doses in PC-3M-Pro4 cells in vitro. **B.** Contour plot of this drug combination showing high Bliss synergy scores. **C, D.** Representative confocal microscopy images of 3D PC-3M-Pro4 spheroids expressing an apoptosis reporter (Flip:GFP-T2A-mCherry) after 12 days of treatment with different Rapalink-1, SBI concentrations, or its combination. The fluorescence intensity ratio of Flip:GFP and mCherry was used to calculate cell death rate. **E.** PC-3M-Pro4 spheroid apoptosis kinetics measured as mean Flip:GFP fluorescence/mCherry fluorescence in cells over 12 days of spheroid culture with drug treatment. **E.** PC3-Pro4 spheroid volume change kinetics measured as the volume of cells over 12 days of spheroid culture with drug treatment.

### 3.5. The synergistic effect of Rapalink-1 and SBI effectively eliminates tumor cells in xenografts and in LAPC9-derived organoids

We next assessed the antitumoral efficacy of the combined treatment in zebrafish xenografts. Treatment with SBI alone up to 50 μM or in combination with 1 μM/nl of Rapalink-1 demonstrated low toxicity, with more than 90% of the larvae surviving the treatment during the experimental timeline (Supplementary **Fig. S7A** and **S7B**). After determining tolerance doses of the treatments, we engrafted PC-3M-Pro4 and PC3 into zebrafish, followed by intravenous injection of the compounds at 4 and 24 hours after cancer cell engraftment. Cancer burden at the metastatic site was measured at 6dpi, showing that the combined treatments had stronger inhibitory effect on the tumor growth than the single treatments (**Fig. 5A, 5B, S8A-G**). At the highest concentration, SBI (10 μM) only caused mild effects, inhibiting 8.26% ± 10.8 of the PC-3M-Pro4 growth in the metastatic foci. Rapalink-1 (1 μM) treatment increased the inhibition to 56.42% ± 9.56 of tumor growth, but the highest effect was found with the combination of both drugs, with a 97.69% ± 1.47 of tumor growth inhibition *in vivo.* Consistent to the *in vitro* experiments, we found a strong synergetic effect of the combined treatment (**Fig. 5C**). Next, we tested impact of the treatments on cell apoptosis *in vivo*. Cancer cells bearing the apoptosis reporter were engrafted into zebrafish and treated with the drug combination. In line with the findings *in vitro*, fluorescence indicative of apoptosis was significantly enhanced when the engrafted fish were co-treated with SBI and Rapalink-1 (**Fig. 5D, 5E**). Collectively, our data suggest that the combination treatment of Rapalink-1 and SBI has a synergistic and strong inhibitory effect on survival and growth of the prostate cancer cells during metastasis onset in a xenograft animal model.

**Figure 5.**
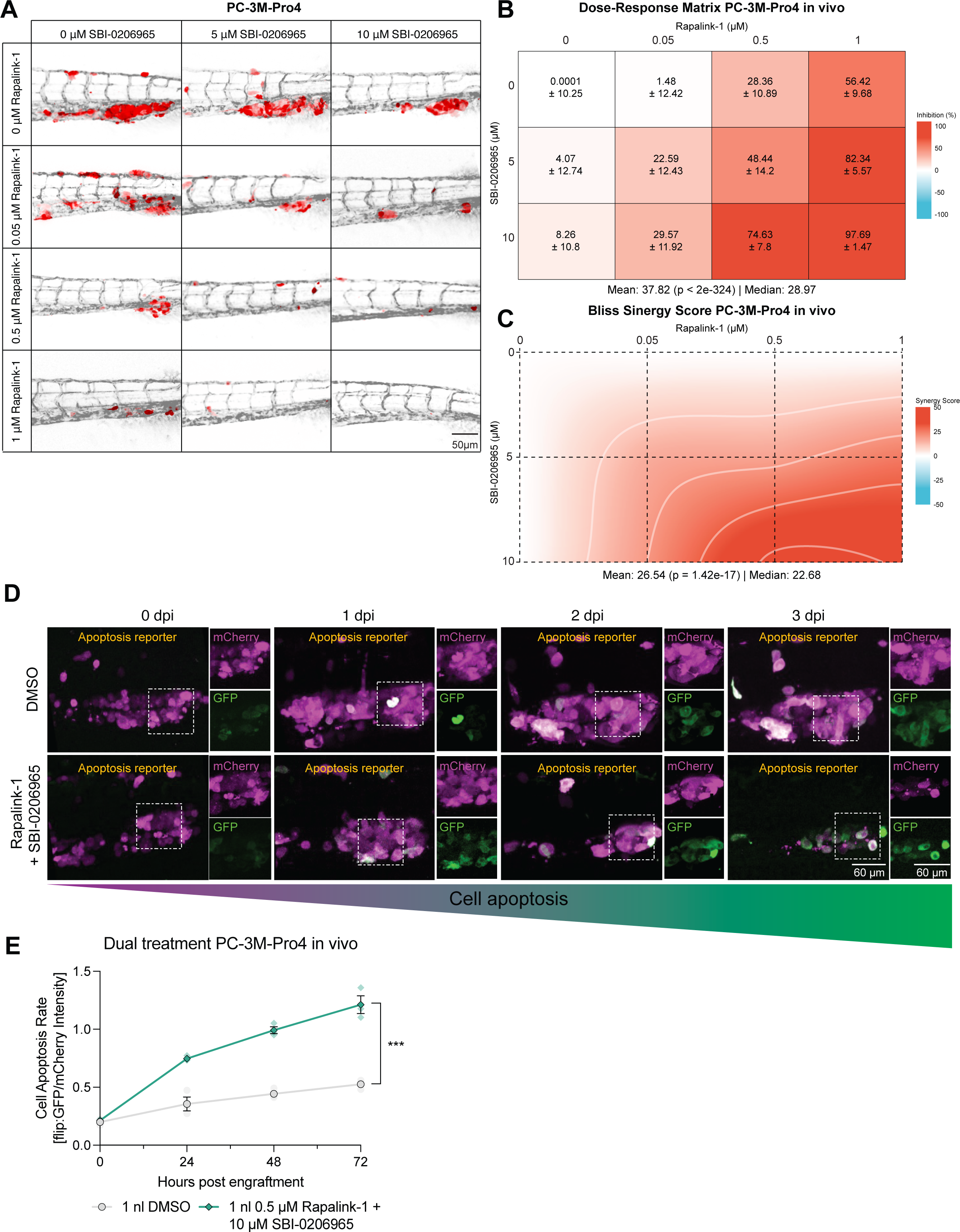
The synergistic effects of Rapalink-1 and SBI are highly effective in suppressing tumor burden in zebrafish. **A.** Representative images of the metastatic foci of PC-3M-Pro4 cells expressing tdTomato at 6 dpi in zebrafish treated with different concentrations of Rapalink-1, SBI, or its combination (n = 15). **B.** Dose-Response matrix of metastatic growth inhibition of different Rapalink-1 and SBI doses in PC-3M-Pro4 cells in vivo. **C.** Contour plot of Bliss synergy score of this drug combination showing high Bliss synergy scores in vivo. **D.** Representative images of PC-3M-Pro4 with stable expression of apoptosis reporter engrafted into zebrafish and treated with DMSO or a combination of 0.5μM of Rapalink-1 and 5μM of SBI at 0 h, 24 h, 48 h and 72 h after administration. **E.** The fluorescence intensity ratio of Flip:GFP and mCherry was used to calculate cell death rate (n = 3).

We then assessed the antitumoral efficacy of the combined treatment using patient-derived LAPC9 organoids [49, 55]. We first confirmed the occurrence of mTOR-AMPK conversion in the organoids in nutrient deficiency. Organoids cultured medium supplemented with 2.5% FCS displayed enhanced AMPK phosphorylation and reduced mTOR phosphorylation, as indicated by immunofluorescence (**Fig. 6A-D**). In order to test whether AMPK could be directly activated by chemical inhibition of mTOR, we transduced the organoids with the AMPK FRET biosensor and treated them with Rapalink-1 (**Fig. 6E**). The FRET signal reflective of AMPK activity was rapidly elevated within 6 hours of Rapalink-1 exposure (**Fig. 6F, 6G** and **Supplementary video 3**). We finally investigated the effects of targeting of AMPK and/or mTOR on organoid growth. Organoids were treated with SBI, Rapalink-1 or combination of both at serial doses. As with previous results, SBI alone exerted very low inhibitory effects, up to 13.95% ± 2.22 of LAPC9 growth inhibition, compared to a 27.1% ± 0.32 inhibition with 0.5 μM of Rapalink-1. However, congruent with previous results, the combined therapy had a synergistic effect up to 82.11% ± 0.74 of LAPC9 growth inhibition (**Fig. 6H, 6I, S9A-D**). Overall, our study highlights that co-targeting of AMPK and mTOR using SBI and Rapalink-1 exerts synergistic antitumoral activity *in vitro*, during metastatic onset *in vivo*, and in LAPC9 patient-derived organoids.

**Figure 6.**
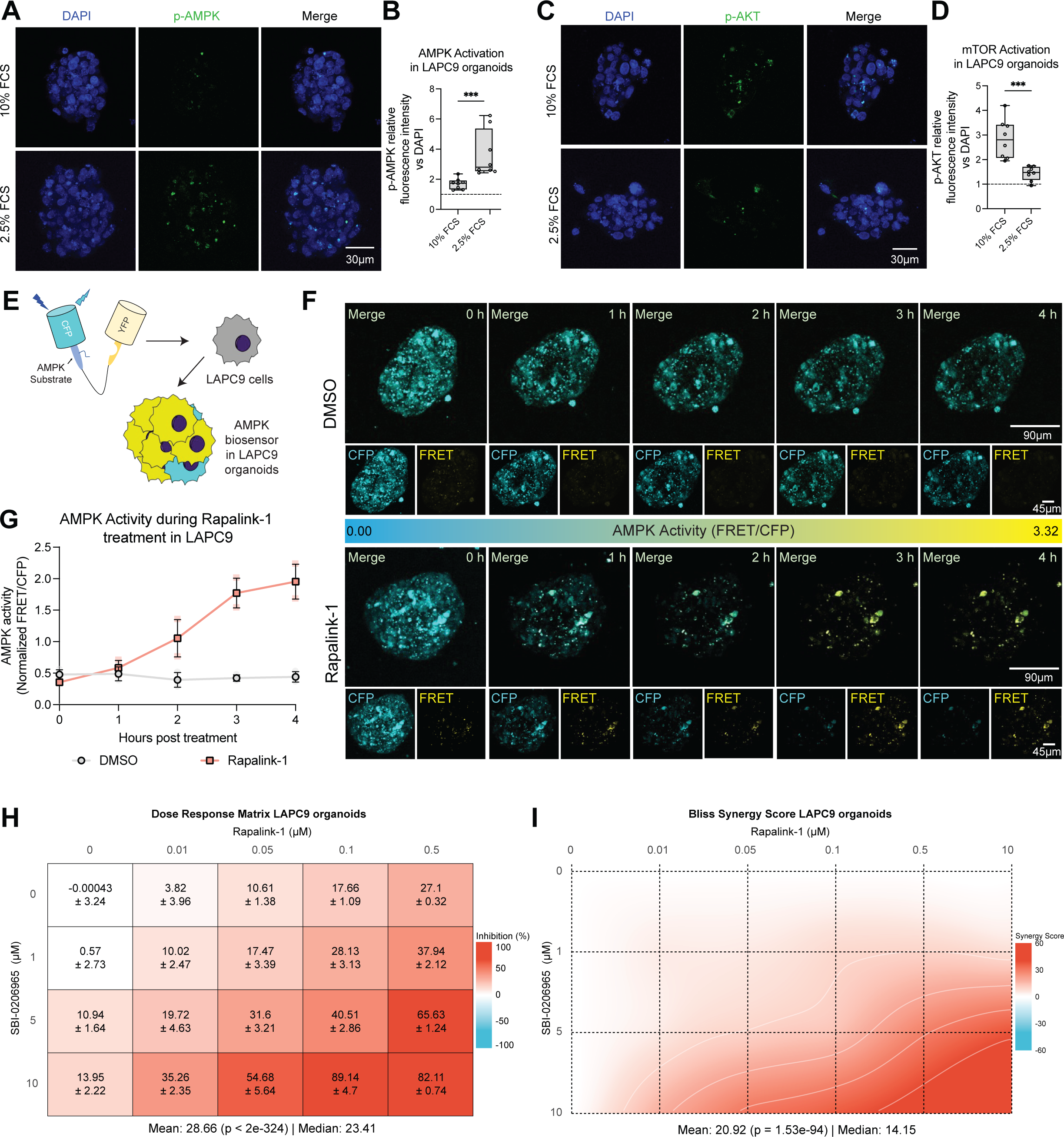
The synergistic effects of Rapalink-1 and SBI are highly effective in suppressing tumor growth in LAPC9 organoids. **A, B.** Immunohistochemistry of p-AMPK in prostate patient-derived organoids LAPC9 cultured on 10% and 2.5% FCS demonstrates AMPK activation under metabolic stress. **C, D.** Immunohistochemistry of p-AMPK in prostate patient-derived organoids LAPC9 cultured on 10% and 2.5% FCS lower mTOR activation under metabolic stress. **E.** Schematic representation of the construction of AMPK biosensor LAPC9 organoids using the AMPK kinase FRET construct. **F, G.** Representative images and quantification of AMPK activity as determined by the AMPK biosensor FRET levels (ratio of FRET, yellow / CFP, blue) shows higher AMPK activity in LAPC9 treated with 0.5 μM of Rapalink-1. **H.** Dose-Response matrix of tumor growth inhibition of different Rapalink-1 and SBI doses in LAPC9 organoids. **I.** Contour plot of Bliss synergy score of this drug combination showing high Bliss synergy scores in LAPC9 organoids.

## 4. Discussion

In the present study, we reveal a critical role of AMPK signaling in regulating prostate cancer cell survival during metastasis. We find that at metastatic onset, AMPK is activated in the disseminated cancer cells concomitant with deactivation of the cell cycle regulator mTOR. This so-called mTOR-AMPK conversion serves as a stress-coping machinery protecting the cancer cells from stress-induced cell death at metastatic onset and thus promotes metastasis.

AMPK has been long considered a tumor suppressor in PCa due to its capacity of inhibiting cell division by metabolism reprogramming and suppression of mTOR signaling [56]. When AMPK was constitutively activated in PTEN-null PCa mouse models, both primary and metastatic tumor growth were significantly suppressed [57]. However, due to a lack of tools for real-time cell signaling analysis *in vivo*, the putative protective role of AMPK in survival of the disseminated cancer cells is largely underestimated in such studies. Using the transparent zebrafish xenografts, we were able to profile the kinetic activities of AMPK and mTOR signaling in the disseminated PCa cells. We observed that the mTOR-AMPK conversion occurred in the cells in circulation, and was reversed when metastatic outgrowth was already established, supporting that active AMPK is required for onset of metastasis but not for metastatic outgrowth. In agreement with this, knockdowns of AMPK induced massive cell death in circulating tumor cells and thus suppressed metastasis potential, indicating the transient activation of AMPK is indeed critically needed for establishment of metastatic onset. It is still unclear how the disseminated cancer cells are protected by active AMPK signaling. In metastasis, upregulation of AMPK is associated with dysregulation of gene signatures indicative of metabolism, suggesting the AMPK activation is associated with metabolic challenging [58]. Consistent with this, when we cultured cancer cells in nutrient deficiency, AMPK was activated to sustain cancer cell survival and growth. In addition, AMPK may be able to protect the circulating tumor cells from other types of stress caused by detachment from extracellular matrix [59] and/or accumulation of reactive oxygen species, through metabolism reprogramming or activation of anti-oxidant pathways (e.g. FOXO signaling) as suggested in other type of tumors [60]. Of interest, AMPK can be directly activated by the loss of mTOR. This dynamic cooperation between AMPK and mTOR is thought to play important role in cell state decision (survival or growth) in distinct contexts.

ACK/PI3K/mTOR signaling is a major driver of PCa progression [61] and has long been the target of chemotherapeutic treatments in the clinic. However, the treatment efficacy of mTOR inhibitors is relatively low [62, 63]. Having shown that AMPK can be directly activated to protect the cells from mTOR inhibition, we hypothesized that the hyperactivation of AMPK can be a cause of the mTOR inhibitor resistance, and co-targeting of both AMPK and mTOR can have strong synergetic inhibitory effect on cancer cell growth. This thought is supported by the fact that transduction of constitutively activated AMPK in PCa cells completely diminished the inhibitory effect of the mTOR inhibitor Rapalink-1, and the knockdown of AMPK, sensitizes the cells to the mTOR inhibition. As such, the anti-cancer efficacy of the combination of the Rapalink-1 and the AMPK inhibitors was tested in 2D/3D cell cultures, zebrafish xenograft model and LAPC9 organoid model. In all tested models, the combined treatment showed strong synergetic inhibitory effects on the PCa cells, indicating a high clinical potential of co-targeting of both AMPK and mTOR.

Although in the present study the inhibitory effect of the co-targeting of AMPK and mTOR has been evaluated in multiple PCa cell lines –including one organoid system–, it would be interesting to further test efficacy of the combined treatment on PCa cell lines/organoids with strong resistance to mTOR inhibition.

Overall, in this study we identified a stress-coping machinery governed by AMPK-mTOR signaling conversion that is critically required for PCa metastasis. Co-targeting of both proteins exerts strong synergetic inhibitory effect on metastatic PCa cells and can be considered as a therapeutic approach against the disease.

## Acknowledgements

We thank Prof. Marianna Kruithof-de Julio (Department for Biomedical Research, Bern University) and Dr. Sylvia Le Dévédec for providing experimental material. Prof. Rob Hoeben and Martijn Rabelink (Department of Cell Biology, LUMC) for supplying lentiviral shRNA vectors (Sigma-Aldrich). G. Zhao gratefully acknowledges the China Scholarship Council (CSC) for personal grants (No. 201906340159). The present work was supported by a personalized medicine grant from Alpe D’HuZes (AdH)/KWF PROPER entitled “Near-patient prostate cancer models for the assessment of disease prognosis and therapy” (UL2014-7058) to B. E. Snaar-Jagalska.

---

**Figure S1.**
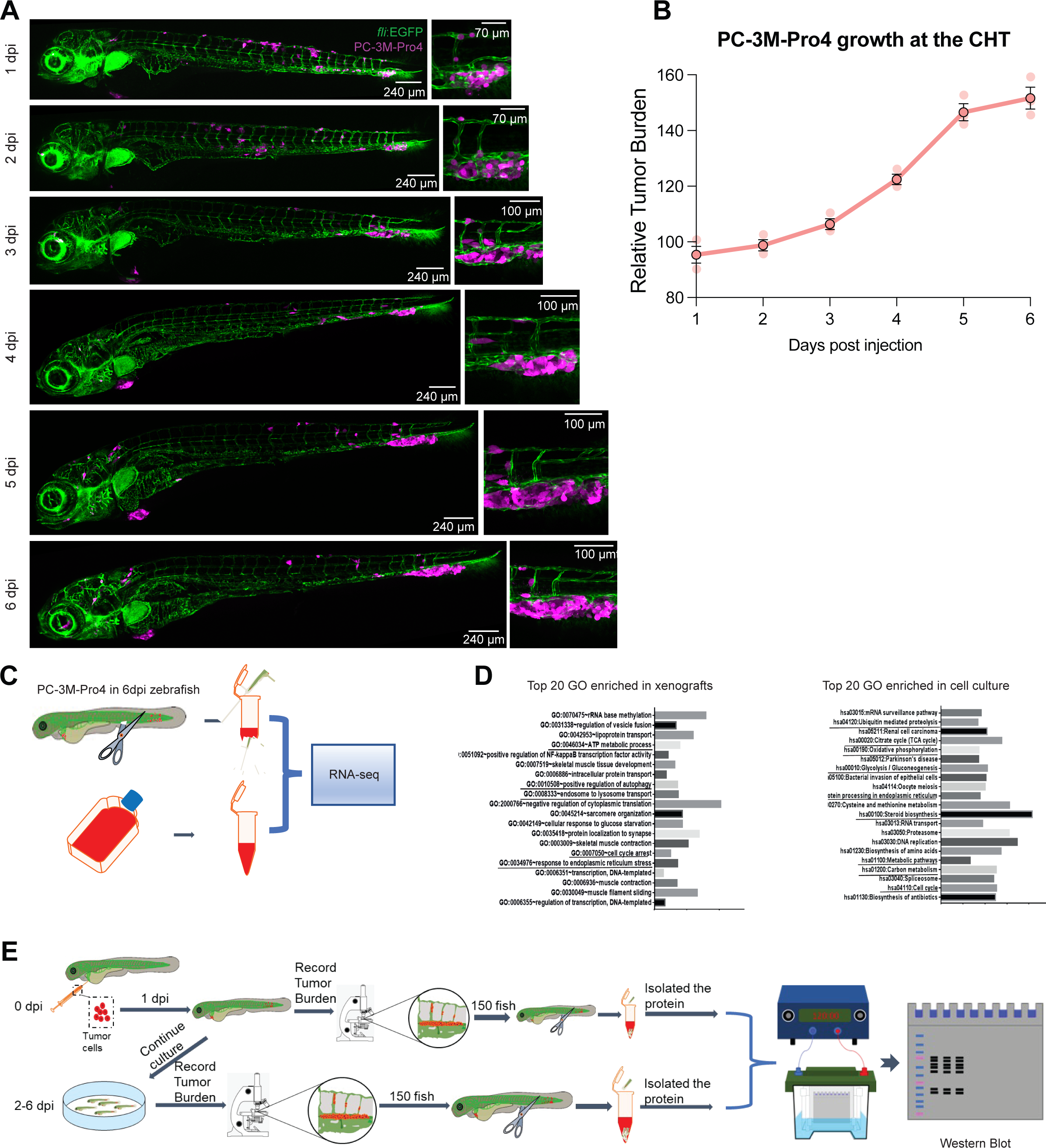
Prostate cancer cells in circulation and RNA-seq data analysis. **A.** Representative images of PC-3M-Pro4 cancer cells expressing tdTomato (magenta) in fli:gfp (green) zebrafish xenografts from 1 to 6 dpi. **B.** Tumor burden can be quantified using the cancer cell line integrated fluorescence intensity. **C.** Schematic indication of RNA-seq analysis of PC-3M-Pro4 cultured in vitro and 6 dpi after xenografting in zebrafish embryos. **D.** Gene ontology analysis was performed using DAVID. The top 20 KEGG pathways enriched in metastasis and in cell culture are displayed (pLJ<LJ0.05). **E.** Schematic diagram of the purification of tumor cell protein samples from zebrafish xenografts for western blot detection.

**Figure S2.**
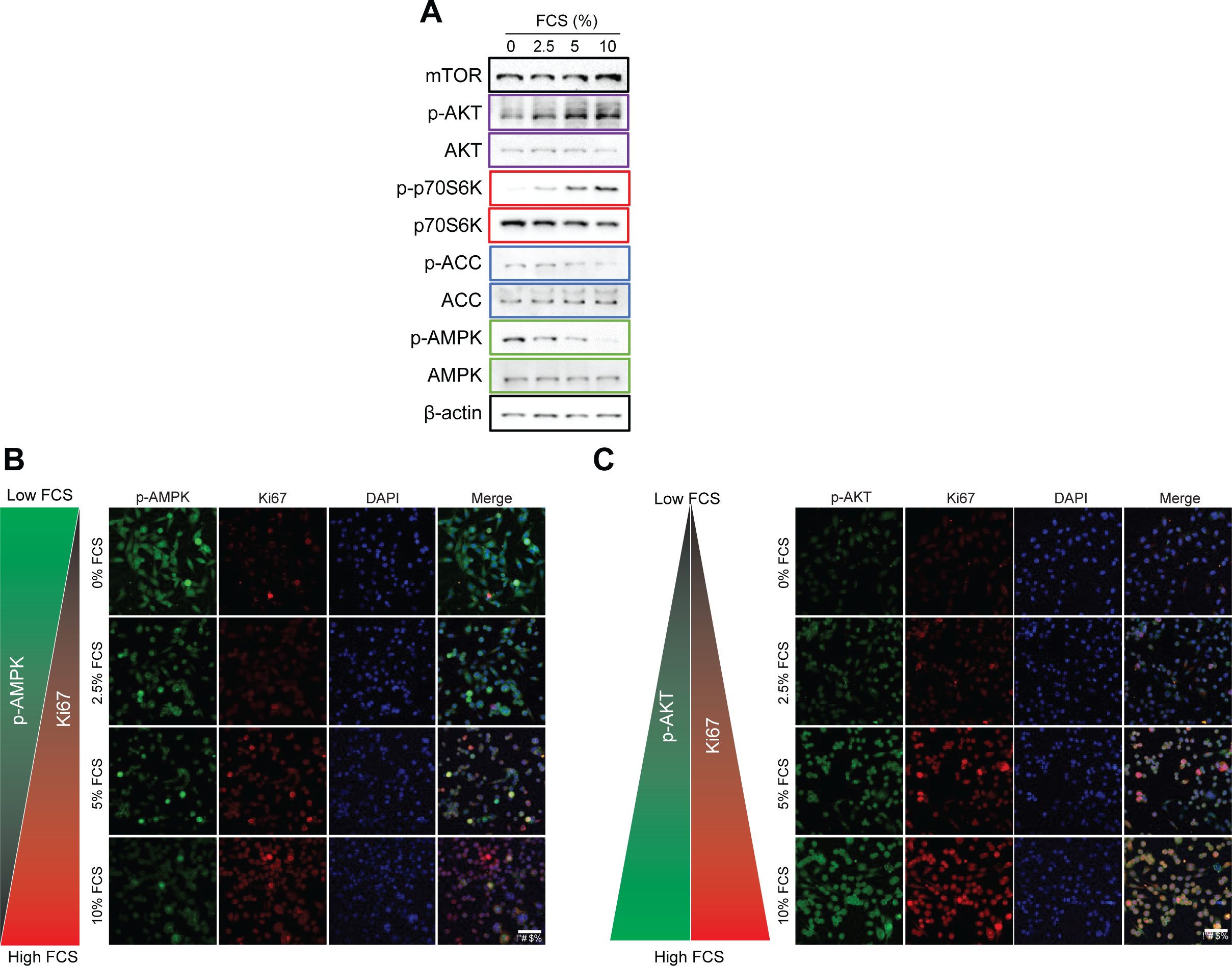
FCS starvation induces activation of AMPK and inhibition of mTOR and thus mediates cell proliferation *in vitro*. **A.** Western blot of phosphorylation levels of proteins downstream of mTOR and AMPK activation in prostate cancer cells in culture at %, 2.5%, 5%, 10% FCS present in the medium. **B, C.** PC-3M-Pro4 cells showed increased AMPK (p-AMPK) and proliferation (Ki67) and mTOR activation (p-AKT) at lower FCS percentage in the medium as demonstrated via immunohistochemistry.

**Figure S3.**
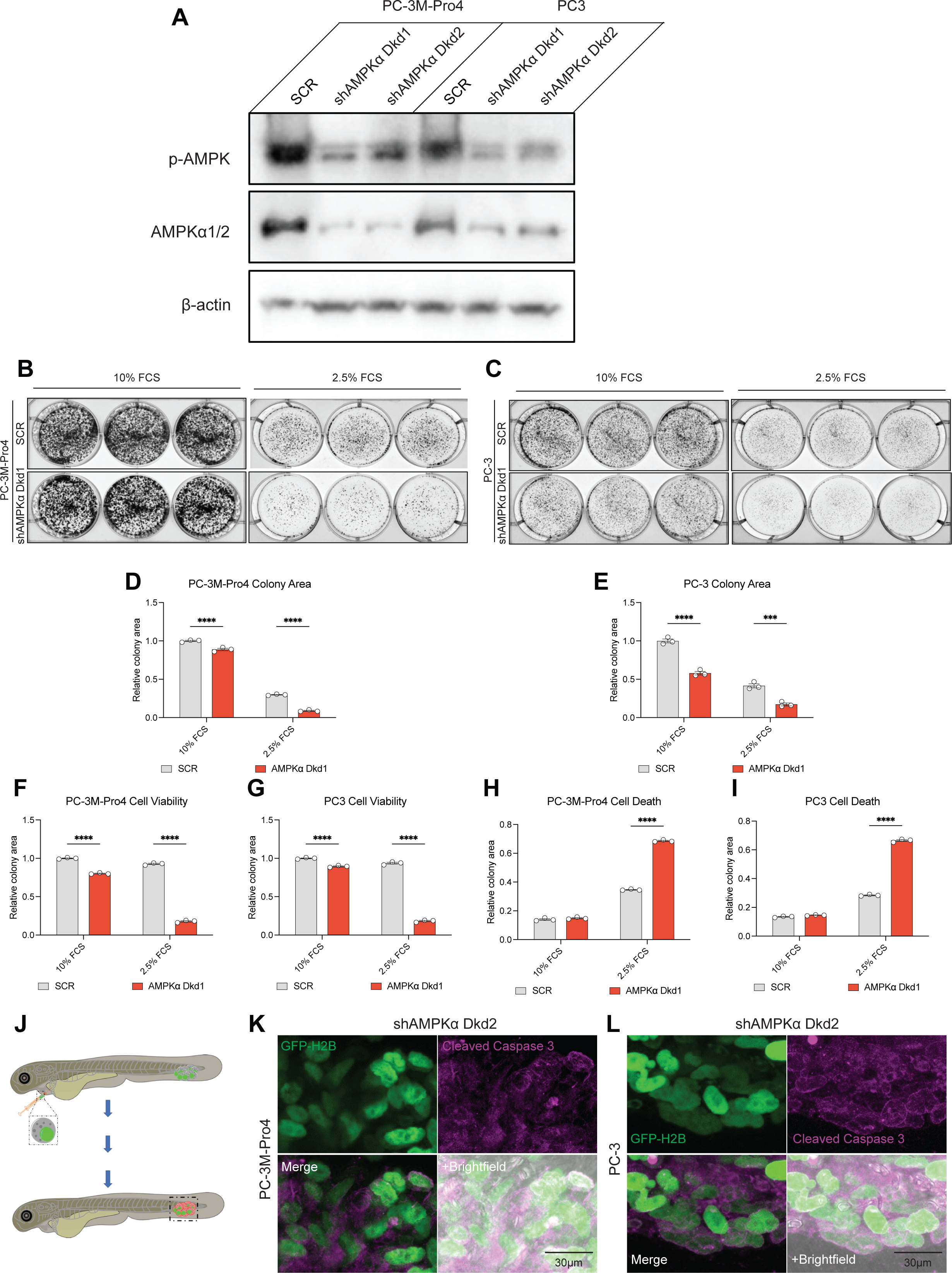
AMPK knockdown inhibits the survival and proliferation of PC-3M-Pro4 and PC-3 cancer cells during starvation. **A.** Transduction with lentivirus encapsulating shRNA targeting AMPKα1 and AMPKα2 knocked down these proteins expression as detected by western blot. **B, D.** Colony assay was used to detect the proliferative ability of PC-3M-Pro4 with AMPKα knocked down at 10% and 2.5% FCS in the medium. **C, E.** Colony assay was used to detect the proliferative ability of PC-3 with AMPKα knocked down at 10% and 2.5% FCS in the medium. **F, G.** Wst-1 kit was used to detect the cell viability of PC-3M-Pro4 and PC3 with SCR, or AMPKα Dkd at 2.5% and 10% FCS. **H, I.** LDH kit was used to detect the cell viability of PC-3M-Pro4 and PC3 with SCR, or AMPKα Dkd at 2.5% and 10% FCS **F**. Scheme of whole amount tumor cell immunostaining in zebrafish. **J-L.** Immunostaining of cleaved-caspase 3 antibody (magenta) was used to detect the cell death of PC-3M-Pro4 and PC3 AMPKα Dkd2 in casper zebrafish, whose nucleus is labelled with GFP-H2B (green).

**Figure S4.**
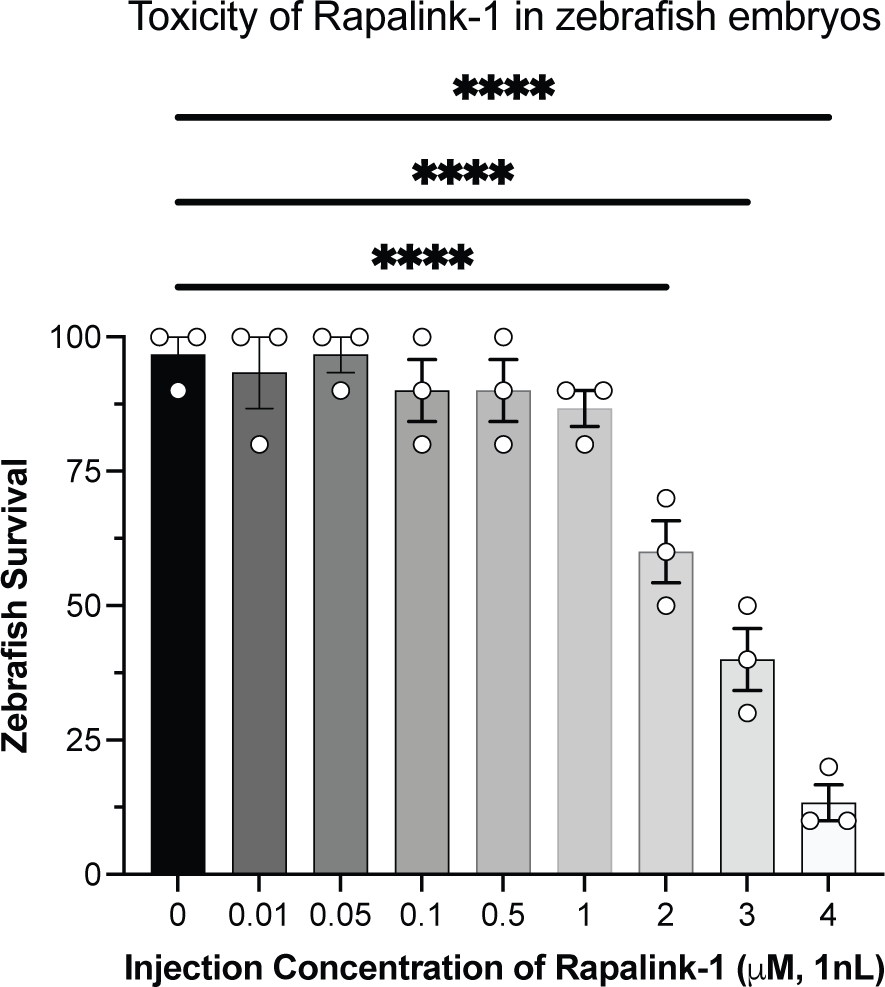
Rapalink-1 is not toxic at effective concentrations in zebrafish embryos. 1 nl of different concentrations of Rapalink-1 were injected into zebrafish by IV administration at 2 and 3dpf and tested for toxicity up to 8 dpf. Rapalink-1 was only toxic at injected concentrations of more than 2μM (1nl).

**Figure S5.**
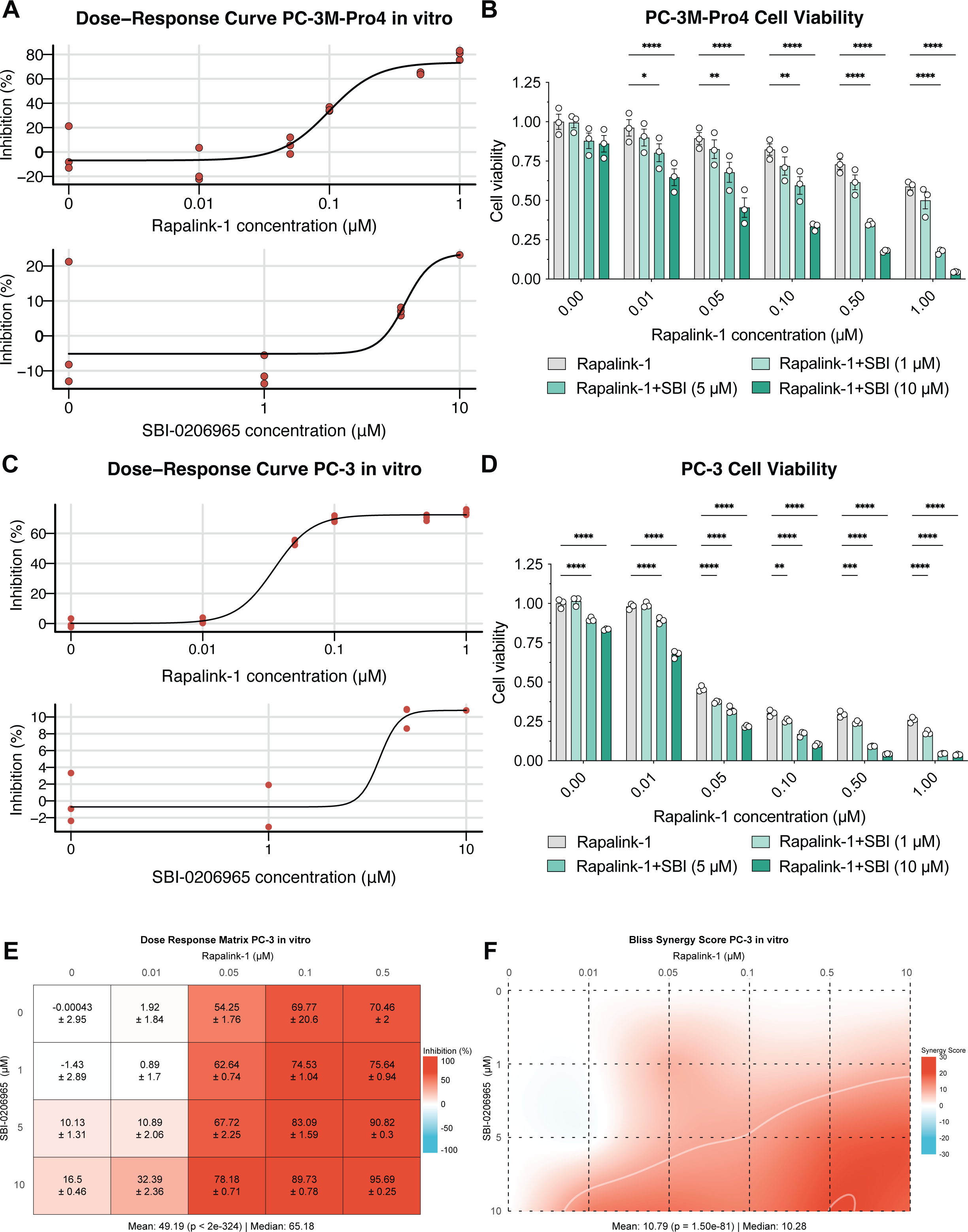
Synergistic use of mTOR and AMPK inhibitors is highly effective in tumor suppression in vitro. **A.** Dose-response growth inhibition curve of Rapalink-1 and SBI on PC-3M-Pro4 cells in vitro. **B.** WST-1 was used to detect the cell viability of PC-3M-Pro4 at different concentrations of Rapalink-1, SBI and the combination of the two drugs. Compared with single drug treatment, the drug combination achieved a cell viability. **C.** Dose-response growth inhibition curve of Rapalink-1 and SBI on PC-3 cells in vitro. **D.** WST-1 was used to detect the cell viability of PC-3 at different concentrations of Rapalink-1, SBI and the combination of the two drugs. Compared with single drug treatment, the drug combination achieved a cell viability. **E.** Dose-Response matrix of cell growth inhibition of different Rapalink-1 and SBI doses in PC-3 cancer cells in vitro. **F.** Contour plot of this drug combination showing high Bliss synergy scores.

**Figure S6.**
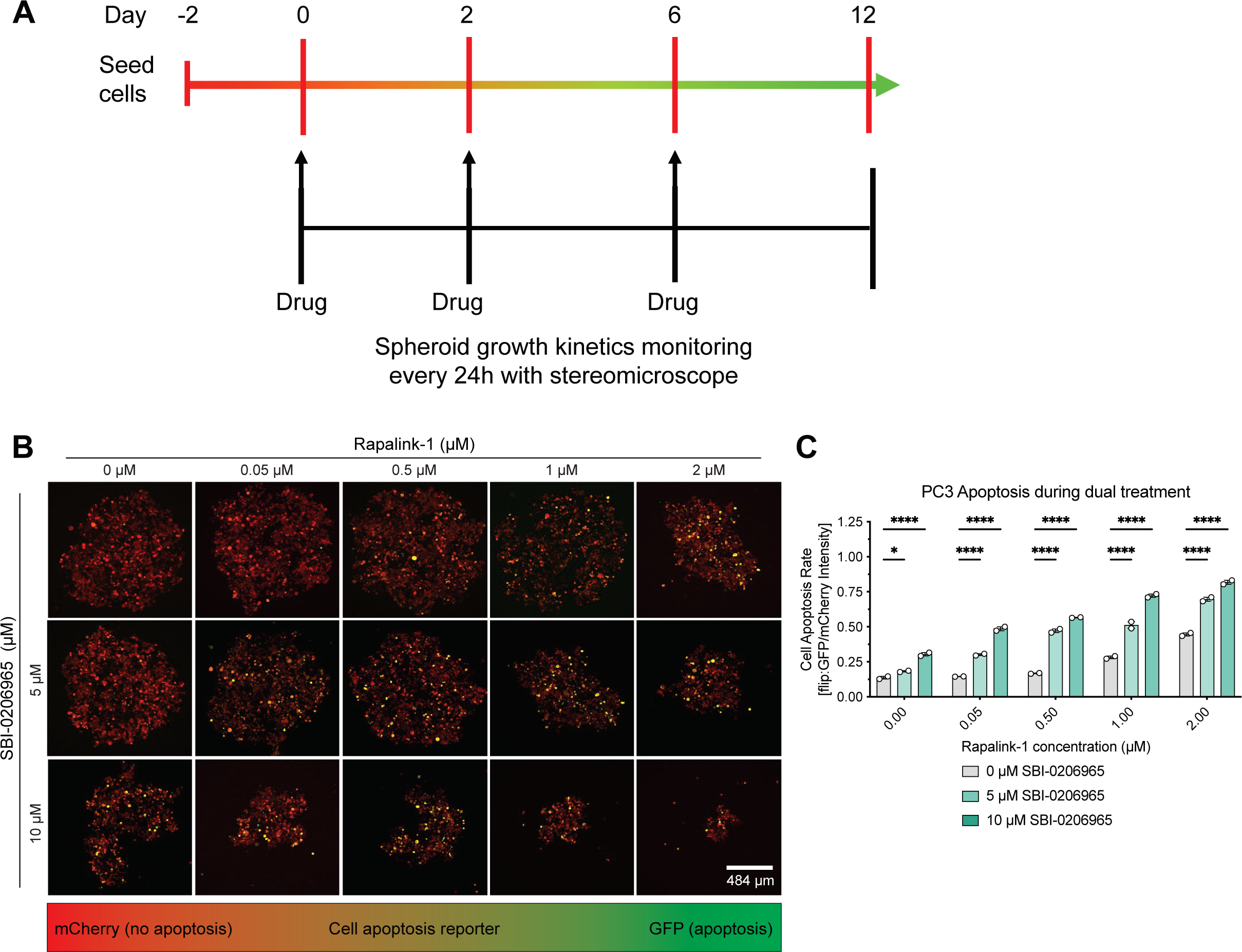
Synergistic use of mTOR and AMPK inhibitors is a highly effective tumor suppressor in 3D cultures. **A.** Schematic representation of the 3D drug efficacy experiment over 12 days. **B.** Representative confocal microscopy images of 3D PC-3 spheroids expressing an apoptosis reporter (Flip:GFP-T2A-mCherry) after 12 days of treatment with different Rapalink-1, SBI concentrations, or its combination. **C.** The fluorescence intensity ratio of Flip:GFP and mCherry demonstrates a concentration-dependent apoptosis rate in single and in combination treatments.

**Figure S7.**
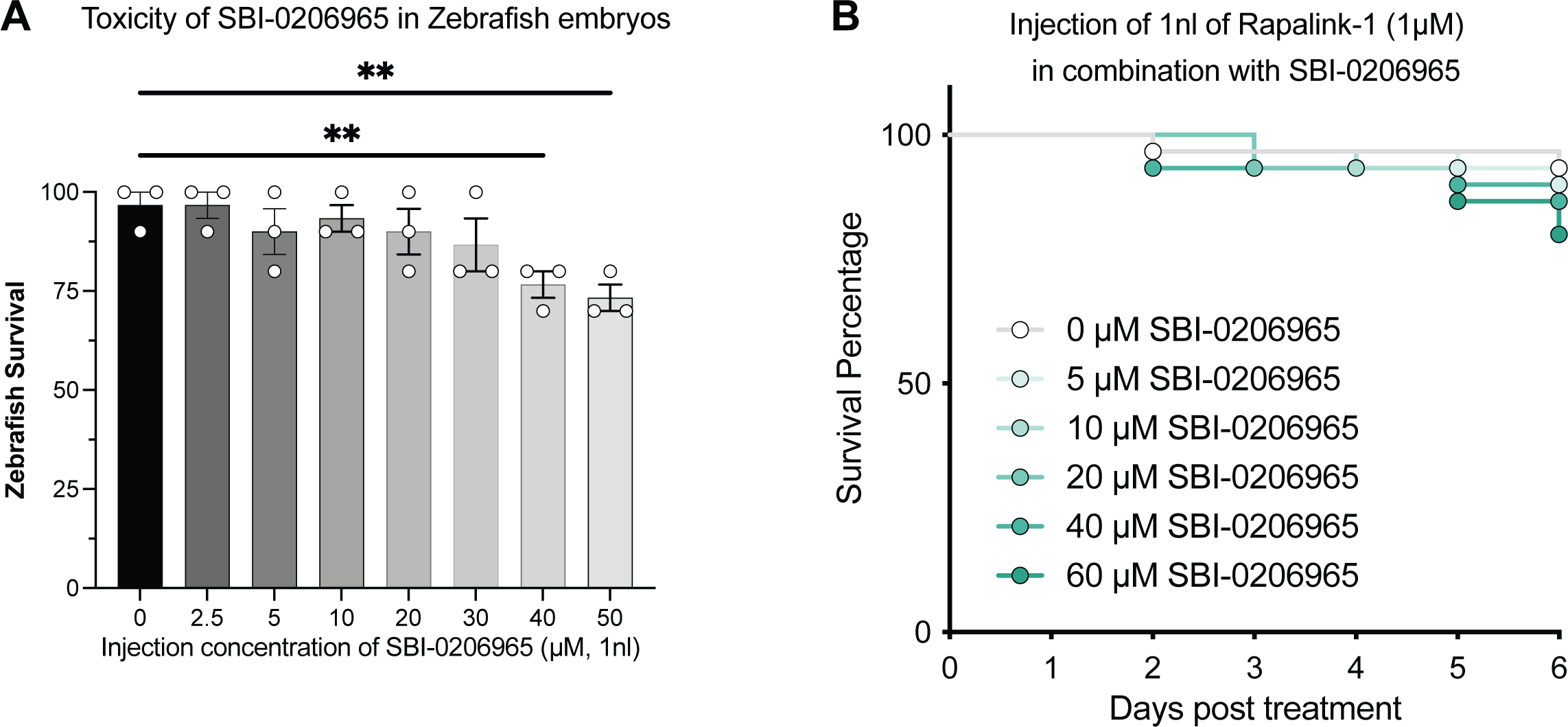
SBI is not toxic at effective concentrations in zebrafish embryos. **A.** 1 nl of different concentrations of SBI were injected into zebrafish by IV administration at 2 and 3dpf and tested for toxicity up to 8 dpf. SBI was only toxic at injected concentrations of more than 50μM (1nl). **B.** To test toxicity to the combination treatment, a fixed dose of Rapalink-1 (1 μM) was injected in combination of increasing concentrations of SBI, which showed low toxicity at concentrations lower than 40 μM.

**Figure S8.**
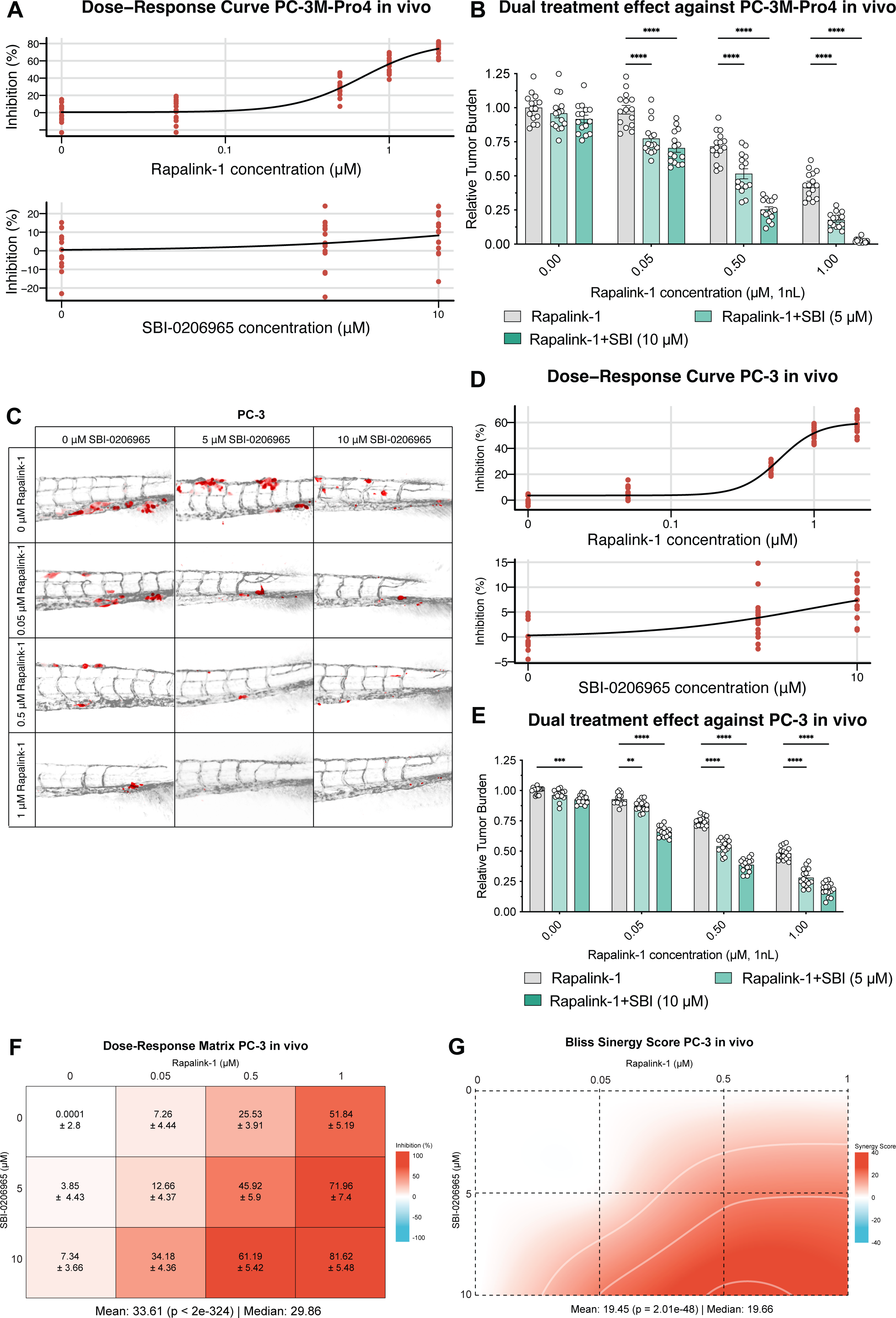
Synergy of Rapalink-1 and SBI is highly effective in tumor suppression in vivo. **A.** Dose-response growth inhibition curve of Rapalink-1 and SBI on PC-3M-Pro4 cells in zebrafish xenografts. **B.** Combination treatment of Rapalink-1 and SBI has a significantly higher antitumoral effect than monotherapy in PC-3M-Pro4 xenografts expressing tdTomato as quantified by integrated fluorescence intensity (n = 15 per group). **C.** Representative images of the metastatic foci of PC-3 cells expressing tdTomato at 6 dpi in zebrafish treated with different concentrations of Rapalink-1, SBI, or its combination (n = 15). **D.** Dose-response growth inhibition curve of Rapalink-1 and SBI on PC-3 cells in zebrafish xenografts. **E.** Combination treatment of Rapalink-1 and SBI has a significantly higher antitumoral effect than monotherapy in PC-3M-Pro4 xenografts expressing tdTomato as quantified by integrated fluorescence intensity (n = 15 per group). **F.** Dose-Response matrix of cell growth inhibition of different Rapalink-1 and SBI doses in PC-3 cancer cells in zebrafish xenografts. **G.** Contour plot of this drug combination showing high Bliss synergy scores.

**Figure S9.**
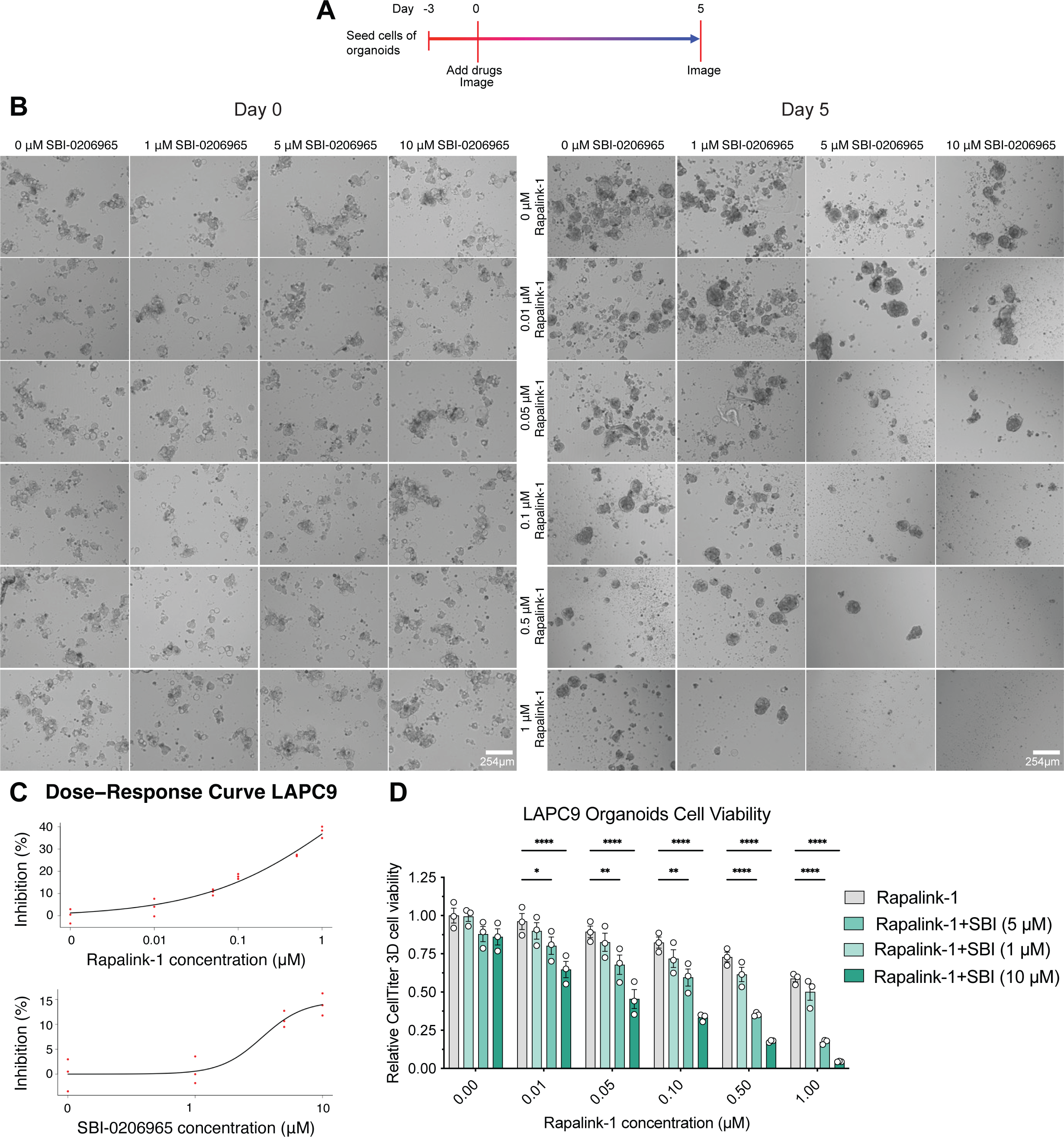
3D titer cell viability detects synergistic effect of Rapalink-1 and SBI in LAPC9 organoids. **A.** Schematic representation of the LAPC9 organoid culture and drug treatment. **B.** Representative images of LAPC9 organoids treated with SBI and Rapalink-1 at different concentrations for 5 days showed that the combination treatment has a stronger tumor inhibitory effect. **C.** Dose-response growth inhibition curve of Rapalink-1 and SBI on LAPC9 organoids. **D.** LAPC9 organoids were treated with Rapalink-1, SBI, or its combination and cell viability was detected at 72h using a 3D titer cell viability assay, showing a strong combinatorial effect.

## Notes

### Competing Interest Statement

The authors have declared no competing interest.

